# Development and validation of KASP markers for a novel powdery mildew resistance gene in wheat using BSR-Seq analysis

**DOI:** 10.1101/2025.04.12.648525

**Authors:** Ramandeep Kaur, Vikrant Tyagi, Raman Dhariwal, Imran Sheikh, Harcharan S. Dhaliwal, M. Sivasamy, Thamaraikannan Sivakumar, Sundeep Kumar, Vikas Kumar Ravat, Neeraj K. Vasistha

## Abstract

Wheat powdery mildew occurs as a devastating infection caused by the pathogenic fungi *Blumeeria graminis f. sp. tritici (Bgt).* The management of disease becomes much more effective when host resistance methods are employed. The WL711-2U/2B line served as the introgressed line to identify genes linked with powdery mildew resistance. This allowed us to look into the potential resistance components in response to powdery mildew infection. The resistance status of the line towards powdery mildew developed from a single dominant gene named *PmAT.* Bulked segregant RNA-Seq (BSR-Seq) together with previous QTL-Seq SNP markers helped identify strong candidate regions in chromosome arm 2B extending from 531114848 bp to 530235618 bp. The LOD score from this region reached 72.4 with a PVE% of 64.7, thus proving its main role in resistance. The physical position was subsequently locked up, which was different from the previously identified genes on the same chromosome arm in its position, suggesting that it is most likely a new *Pm gene*. The QTL interval contained five potential genes that we identified during this research. The five candidate genes directs the protein production, which performs functions linked to the pentatricopeptide repeat family as well as B-box-type zinc finger domain, P-loop-containing nucleoside triphosphate hydrolase, and plant peroxidase.

**Highlight:** The identification of a new wheat powdery mildew resistance gene (*PmAT*) using a combination of BSR-Seq and QTL-Seq analysis, which located five candidate genes specifically on chromosome 2B.

## Introduction

Bread wheat (*Triticum aestivum* L. em Thell.) is serving as important source of nutrition and most significant staple cereal crop for around one-third of the world’s population including India (Singh et al., 2024). Global wheat production is at a record 787.70 million metric tons (MMT) in 2023-24 and the top five wheat-producing countries in the world for the 2023-2024 marketing year are China, European Union, India, Russia and United States (USDA, 2024). Wheat grain production rates in Indian farmers’ fields have increased by more than 1.8 percent annually per hectare during the last decade thereby exceeding global averages by significant margins (1.3 percent). The introduction of improved wheat varieties to farmers, due to greater policies and approaches that speed up seed multiplication, along with greater involvement of private seed producers has accelerated significantly (Govindan et al., 2023). Accordingly, upto 70% increase in wheat production may help to meet global demand of food by 2050 (Nelson et al., 2010). The average wheat yield has stagnated by 40% during recent years, which demonstrates the current production rate is inadequate for future food security. Powdery mildew disease, together with multiple abiotic and biotic factors, has negatively affected wheat production levels across the entire world.

Wheat faces severe damage from the disease known as powdery mildew. The disease exists naturally in regions with cool temperatures and wet conditions. The study conducted by Basandrai and Basandrai (2017) in Himachal Pradesh, India, found that powdery mildew infection causes damage ranging from 13-34% under low to moderate intensity, however, this damage can increase up to 50% under more severe conditions. Exploiting enhanced breeding techniques facilitates developing novel and numerous resistance genes by using wild relatives as a primary source of resistance, but sometimes it creates problems in wheat breeding by linkage drag and other complications as well (Yumurtaci, 2015). Although many powdery mildew resistance genes have been identified worldwide, only a limited number are currently used in commercial applications. Reports of powdery mildew outbreaks in various countries indicate that the races exhibit lower virulence frequencies against newly identified resistance genes. Except for *Pm35*, the discovered powdery mildew isolates were largely avirulent to the recently disclosed powdery mildew genes (*Pm25–Pm53*), just like in South Africa. Generally, resistance to all pathogen races is conferred by either none or very few genes. Because of this, short-lived and tentative genes for powdery mildew resistance have been found, but their persistence has limited their application in creating resistant cultivars (Golzar et al., 2016; Li et al., 2016).

The development of resistant cultivars that are cultivated to reduce yield losses, boost profitable production, and minimize the risk of linkage drag is aided by breeding techniques. Plants resistant to disease that possess a certain quality is primarily due to the resistant gene, also known as the main R gene. However, the traits associated with resistance are presented measurably and so Quantitative trait loci (QTL) were defined (Pilet-Nayel et al., 2017). Advancements in recent years on new genomic breeding approaches and genotyping technologies based on Next-Generation Sequencing (NGS) have led to a huge role in identifying and introducing powdery mildew resistance traits into commercial cultivars. Over 240 powdery mildew resistance genes/loci and more than 60 genes/alleles have been reported on 21 wheat chromosomes and 18 chromosomes, respectively (Kang et al., 2020). So far, a total of 21 powdery mildew resistance genes or alleles have been successfully cloned which include *Pm1a* (Hewitt et al., 2021), *Pm2* (Sánchez-Martín et al., 2016), *Pm3* (subtypes *Pm3a*, *Pm3b, and Pm3d*) (Srichumpa et al., 2005; Yahiaoui et al., 2004), *Pm4b* (Sánchez-Martín et al., 2021), *Pm5e* (Xie et al., 2020), *Pm8* (Hurni et al., 2014), *Pm12* (Zhu et al., 2023), *Pm13* (Li et al., 2024), *Pm17* (Singh et al., 2018), *Pm21* (He et al., 2018; Xing et al., 2018), *Pm24* (Lu et al., 2020), *Pm36* (Li et al., 2024), *Pm38*/*Yr18*/*Lr34*/*Sr57* (Krattinger et al., 2009), *Pm46*/*Yr46*/*Lr67*/*Sr55* (Moore et al., 2015), *Pm41* (Li et al., 2020), *Pm55* (Lu et al., 2024), *Pm57* (Zhao et al., 2024), *Pm60* (Zou et al., 2018), *Pm64* (Zhang et al., 2021), *Pm69* (Li et al., 2021), and *WTK4* (Gaurav et al., 2022). Among these cloned genes, only seven (*Pm1a, Pm3, Pm4b, Pm5e, Pm24*, *Pm38/Yr18/Lr34/Sr57* and *Pm46/Yr46/Lr67/Sr55*) are associated with hexaploid wheat. Additionally, four genes (*Pm36*, *Pm41*, *Pm64*, and *Pm69*) have been identified in wild emmer wheat (*T. turgidum* L.). The remaining genes are found in various wild relatives of wheat.

Recent progress in genetic resources and the accessibility of efficient genotyping techniques have led to the emergence of new approaches for identifying, locating, and duplicating genes in identifying the resistance regions. Bulked segregant analysis (BSA) as a novel technique helps to identify genetic markers, which linked to expressed genes/QTLs. The variants and gene expression profiling may be detected by RNA sequencing platform and it also generates transcriptome analysis of short sequences. The RNA-seq approach involves relative quantification of gene expression and is also used to identify Single nucleotide polymorphism (SNP). These SNPs working as molecular markers in the further events. The potential of Bulk Segregant Analysis (BSA) with the affluence of RNA sequencing and suitable statistical measures combines to form a new genomic mapping strategy called as Bulked Segregant RNA sequencing (BSR-Seq) analysis, which is highly efficient and low-cost method to rapidly map an R gene (Dakouri et al., 2018; Wang et al., 2017; Wu et al., 2022). This strategy has been utilized to speedily identify genes that provide resistance to wheat stripe rust and powdery mildew (Hu et al., 2019; Wang et al., 2018; Wu et al., 2018; Zhan et al., 2021).

The majority of resistant genes are specific to certain races and is not effective against all races of the powdery mildew pathogen and are employed in wheat breeding to enhance resistance against powdery mildew (Bapela et al., 2023). Additionally, these genes are prone to breakdown due to the introduction of virulent races. It becomes essential to find new resistance sources against powdery mildew. Research at our station enabled Prof. H. S. Dhaliwal to develop various introgression lines of *Aegilops triuncialis* wild wheat relatives for powdery mildew resistance in wheat. The study employs BSR-Seq analysis to locate and authenticate powdery mildew resistant wheat targets that KASP assay will validate. This is the driving force behind our investigation, which uses BSR-Seq to identify and validate resistant genes and genomic loci for powdery mildew in hexaploid wheat through the KASP assay.

## Materials and methods

### Plant materials and Growth conditions

The resistant introgression line WL711-2U/2B was crossed with WL711, a common wheat cultivar that is susceptible to rust and powdery mildew at Eternal University, Baru Sahib, Sirmour, HP, India, to develop a mapping population consisting of 547 F_2_ plants. WL711-2U/2B and WL711, the parental lines, are both stable genotypes. However, WL711-2U/2B and WL711 have different immune and susceptible reactions to powdery mildew infections. For the development of a resistant parental line WL711-2U/2B, the susceptible WL711 was crossed with *Ae. triuncialis* at the Punjab Agricultural University in Ludhiana, Punjab, India, and then the F_1s_ were backcrossed with WL711 (AghaeeLSarbarzeh et al., 2001; Singh et al., 2000). The whole process involved selecting BC_1_F_1_ plants that were resistant to leaf rust, crossing them back with WL711, and then selfing to make BC_3_F_11_ lines (Kuraparthy et al., 2007). Using a mixture of the most virulent local isolates of the *Bgt* pathogen, these plants were subjected to continuous screening for over 15 years under artificial powdery mildew epiphytotic conditions during the adult plant stage at Eternal University, Baru Sahib, Sirmour (Himachal Pradesh), India. One of these lines, which was later designated as WL711-2U/2B, showed resistance to all the powdery mildew pathogen isolates found in the area.

### Pathology assays

Individual F_2_ seeds were sown in 1 meter long rows each with 15 cm plant to plant distance into well prepared soil under a polyhouse along with the five lines of each parent. Susceptible WL711 was sown around the experimental site to increase the inoculum. A 2cm flag leaf section was collected from every F_2_ and its parental lines before infection. A mixed population of the *Bgt* isolates was used in their natural state to create epiphytotic conditions in disease nurseries at the adult plant stage. During the powdery mildew disease development period, the temperature and relative humidity in Baru Sahib generally range between 15L and 22L (max. 30L) and 70 and 90%, respectively. The region is a natural hot spot for powdery mildew occurrence, evidenced by data from the past 15 years (Dr. H. S. Dhaliwal, personal communication).

Disease symptoms were evaluated in 15 days after inoculation and 21 days after inoculation. Plants were classified as resistant or susceptible based upon the presence of sporulation. The plants of the F_2_ population were grouped into 6 bulks comprising either resistant [Bulk - R1 (76 lines), R2 (80), and R3 (82)] or susceptible [Bulk - S1 (42 lines), S2 (45), and S3 (51)] genotypes. Leaf samples were collected from each F_2_ line and two biological replicates of each parental line (5 plants) for RNA extractions.

### RNA Isolation and Library Preparation

For each bulk and parental line, total RNA was isolated using TRIzol reagent (Invitrogen, Paisley, UK) following the manufacturer’s instructions. In total, there were six RNA bulk samples (3 resistant bulks and 3 susceptible bulks) and 4 samples for parental lines WL711-2U/2B and WL711 (resistant parent × 2 biological replicates and susceptible parent × 2 biological replicates). An in-solution DNA digestion was performed using DNA-free™ DNA Removal Kit (Invitrogen, California, USA) following the manufacturer’s protocol to remove DNA contamination from the RNA samples. The absolute mRNA Purification Kit from Agilent Technologies (Santa Clara, CA, United States) was used to retrieve transcripts before cDNA library construction. The cDNA fragments received Illumina paired-end adapters which were followed by barcode sequences during ligation. The pooled libraries were sequenced at Bionivid Technology Pvt. Ltd. (Bengaluru, India) using the Illumina Novaseq 6000 platform (Illumina, San Diego, CA, USA) to generate 151 bp paired-end (PE; 151 × 2) sequence reads for parents and resistant and susceptible bulk samples.

### SNP identification and bulk frequency ratio (BFR) calculation

BSR-Seq reads were stored as FASTQ-compressed files in chunks for parallel processing in the downstream analysis. The quality of the sequencing reads were checked with FastQCv0.10.1. Fastp v0.20.0 (https://github.com/OpenGene/fastp) (Chen et al., 2018) was used for quality filtering and adapter trimming. A quality cutoff of 30 was set for the Phred score and only high-quality reads were retained. The filtered reads were aligned using BWA aligner against the *Triticum aestivum* reference genome (IWGSC CS RefSeq v2.1) using BWA 0.5.9. The alignments were stored as BAM files after sorting and indexing using SAMtools (https://www.htslib.org/). The SAMtools command fixmate fixed read-pairing issues while markdup eliminated potential PCR duplication as well as genome alignment of identical coordinates. Coverage at a single base resolution was computed using SAMtools command mpileup. Variants (SNPs and INDELs) among bulks were called using VarScan v2.3.9 (https://varscan.sourceforge.net/) (Koboldt et al., 2012). To ensure the reliability of variant calls, the VarScan program was configured with specific criteria. These included a minimum read depth of 15 and a minimum of 8 supporting reads per position for a variant call. Additionally, only bases with a quality score of 15 or higher were considered. SnpEff (https://pcingola.github.io/SnpEff/) was used to determine variants functional effect and their annotation.

Bulk frequency ratio (BFR) was calculated from the SNPs detected between de novo assemblies of two parents (WL711-2U/2B and WL711) using VarScan. Specifically, SNPs were identified from the assemblies of both parental lines (WL711-2U/2B and WL711) and bulks using SAMtools and VarScan. Susceptible parent assembly was used as a pseudo-reference to call SNPs from short reads of the bulks. Fisher exact tests using a 0.05 significance level were applied to identify SNPs that showed frequency differences between the two bulks. Then the SNPs found in the bulks were compared to those in the parental lines, ensuring they matched. Quality control measures were applied, only considering SNPs from reads with high base quality scores (Phred 20 or above) and at least 6 reads. Then, potential SNPs were detected by combining those from the bulks with the parental SNPs. Finally, the Bulk Frequency Ratio (BFR) was calculated based on the frequency of the resistant allele in the bulks to pinpoint SNPs linked to powdery mildew resistance, using a BFR value of 15 or more as our threshold.

### Development of KASP markers from identified SNPs

NCBI Genome Data Viewer (https://www.ncbi.nlm.nih.gov/genome/gdv/browser/genome/?id) was used to retrieve 150 bp long DNA sequence around each selected SNP showed polymorphism and association with powdery mildew resistance. PolyMarker pipeline (http://www.polymarker.info/) is used to design the KASP primers from the retrieved sequences. For each SNP, two allele-specific forward primers and one common reverse primer were designed to test their ability to differentiate the resistance and susceptibility. The KASP protocol was modified to visualise the results on the agarose gel in which we added non-plant-based nucleotide sequence (20 nt) in a forward primer out of two. The KASP primers were then verified on the F_2_ population (114 lines). KASP assay was performed in a reaction mixture of 4.0 µL [2.0 µL DNA (5ng/µL), 1.944 µL of 2X KASP mix, and 0.056 µL primer mix] in a 96-well format thermal cycler (Applied Biosystems, Foster City, California, USA) following PCR conditions: hot-start activation at 95°C for 15 min, followed by 10 touchdown cycles (95°C for 20 s then touchdown at 61°C initially and decreasing by 0.6°C per cycle from cycle 2), and then 30 additional cycles of annealing (95°C for 20 s then 55°C for 60 s). The band pattern was visualised in the SYNGENE Gel Documentation system (Cambridge, UK) using 3% agarose gel.

### Microsatellite marker analysis on chromosome 2B

Genomic DNA was extracted from leaf samples collected before infection using the cetyltrimethylammonium bromide (CTAB) method (Stein et al., 2001). Microsatellite markers positioned on wheat chromosome 2B were selected to precisely map the detected powdery mildew resistance gene. The Primer sequences were obtained from the GrainGenes 2.0 (https://graingenes.org/GG3/) and synthesized by Eurofins Genomics India Pvt. Ltd., Bengaluru, India. The markers were first used to test the parents and both bulks. Only the markers polymorphic between the parents and bulks were used to genotype the 114 F_2_ lines selected randomly. The polymerase chain reaction (PCR) amplifications for markers were performed in a total volume of 20.0 μl using 2.0 μl 10× PCR buffer (Takara, Shiga, Japan), 1.2 μl MgCl2 (25 mM), 0.3 μl dNTP (10 mM), 1 μl each of forward and reverse primers (10 mM), 0.25 μl Taq DNA polymerase (5U/ μl) (Takara, Shiga, Japan), 11.25 μl nuclease-free water, and 3 μl DNA template (50 ng/μl) using an S1000 Thermo Cycler (BioRad, California, USA). Using the touchdown PCR amplification method we performed initial denaturation at 94°C for 4 minutes followed by 10 cycles that included denaturation at 94°C for 1 minute besides touchdown annealing from 68°C down to 65°C (each cycle with a 0.8°C decrease) which occurred for 1 minute and extension at 72°C for 1 minute. The next 35 cycles proceeded with denaturing at 94°C for 1 minute followed by annealing at 72°C for 1 minute using a constant temperature between Tm minus 5°C. A final step of the reaction lasted 10 minutes at a heat setting of 72°C. A 3% high-resolution agarose gel prepared in LONZA (Basel, Switzerland) 0.5X TBE buffer with 0.5 μg/ml ethidium bromide at a stock concentration of 10 mg/ml separated the PCR products. Data visualization and documentation of gels occurred through the SYNGENE Gel Documentation System located in Cambridge (UK).

### Development of linkage map using previously and newly identified molecular markers

F_2_ population (110 plants) genotyping data sets belonging to newly developed KASP markers and microsatellite markers were utilized to construct the linkage map utilizing QTL IciMapping v3.2 software (http://www.isbreeding.net). A LOD threshold of 3.0 was applied between adjacent markers, following the method described by (Li et al., 2007). Mapping function Kosambi (Kosambi 2016) was used for estimating genetic distances between markers. For the identification of most significant markers, the single marker-analysis (SMA) algorithm for additive gene effects was utilized. This algorithm is implemented in QTL IciMapping v.3.2 software, which is a valuable tool for determining the contribution of individual markers to the expression of powdery mildew resistant QTLs. Total 10 molecular markers were utilised for the construction of the linkage map. In which seven were SNPs and the remaining three markers were SSRs (**Table 1**).

**Table 1.** List of molecular markers used in the construction of linkage map.

| S. No. | Marker Name | Type of Marker | Method for the Identification of markers |
| --- | --- | --- | --- |
| 1. | *PmR531114848* | SNP | In present study (BSR-Seq analysis) |
| 2. | *PmR532036320* | SNP | In present study (BSR-Seq analysis) |
| 3. | *PmR532202741* | SNP | In present study (BSR-Seq analysis) |
| 4. | *PmR533100430* | SNP | In present study (BSR-Seq analysis) |
| 5. | *PmR533542998* | SNP | In present study (BSR-Seq analysis) |
| 6. | *PmR530235618* | SNP | QTLseq study (Unpublished data) |
| 7. | *PmR530235656* | SNP | QTLseq study (Unpublished data) |
| 8. | *Xbarc349* | SSR | In present study (Validation in parental lines) |
| 9. | *Xgwm526* | SSR | In present study (Validation in parental lines) |
| 10. | *wmc627* | SSR | In present study (Validation in parental lines) |

### Identification of candidate genes

Further discovering candidate genes from the most important QTLs discovered throughout the study. The markers associated with QTLs received sequence details, which were compared to the IWGSC v2.1 wheat genome assembly found in Ensembl to identify candidate genes in designated regions. Our research examined the interval areas within QTL regions to identify highly important genes with defined functions.

## Results

### RNA Sequencing and SNPs discovery

We applied BSR-Seq to detect resistant loci in wheat against the powdery mildew pathogen. We used the Illumina Novaseq 6000 platform to paired-end sequence a total of 10 RNA samples belonging to parents and extreme bulks. The four parental samples consist of RP1 (resistant parent replication 1), RP2 (resistant parent replication 2), and SP1 (susceptible parent replication 1), and SP2 (susceptible parent replication 1), which are susceptible. There are also six extreme phenotypic bulked samples: RB1 (resistant bulk replication 1), RB2 (resistant bulk replication 2), RB3 (resistant bulk replication 3), SB1 (susceptible bulk replication 1), SB2 (susceptible bulk replication 2), and SB3 (susceptible bulk replication 3). RNA sequencing of these samples produced 68399366, 62598101, 66507494, 61736700, 60873158, 52753499, 55105620, 57729151, 56032476, and 69599370 raw reads, respectively (Table 2). After quality control, between 3% and 11% of the raw reads were filtered to get rid of reads from different samples (P-value < 0.05). This process resulted in the mapping of 88.81% to 97.35% of the reads to the reference genome. Following mapping, we merged the BAM files from all three resistant bulks, and similarly, we merged the BAM files from all three susceptible bulks. We merged the BAM files from each parent in a similar manner. Ultimately, we obtained four combined BAM files: RP for the resistant parent, SP for the susceptible parent, RB for the resistant bulk, and SB for the susceptible bulk. SNP calling successfully identified 85,827 high-quality variants (SNPs and indels) between RB and SB bulks. Around 90% of SNPs in each combined sample were of high quality (p value < 0.1) (Fig. 1A). Although SNPs were found on all of the 21 chromosomes but chromosome 2B contained the maximum SNPs (22393) followed by 2A (Table 3, Fig. 2B).

**Figure 1.**
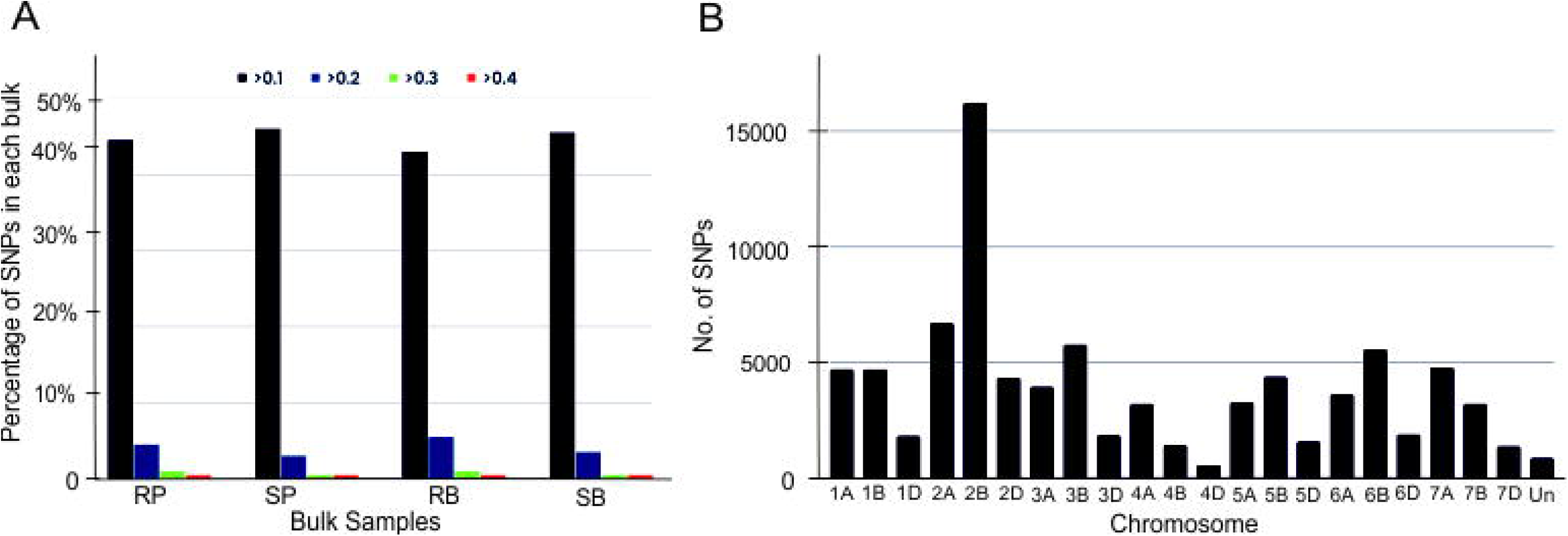
The number of SNPs per 100 base pair (bp) reads and their distribution on the chromosomes. **A)** Percentage of SNPs per 100 bp reads belonging to resistant parent (RP), susceptible parent (SP), resistant bulk (RB) and susceptible bulk (SB), at the p <0.1, <0.2 and <0.3. **B)** Distribution of the polymorphic single-nucleotide polymorphisms (SNPs) on each chromosome analysed using the BSR-Seq.

**Figure 2.**
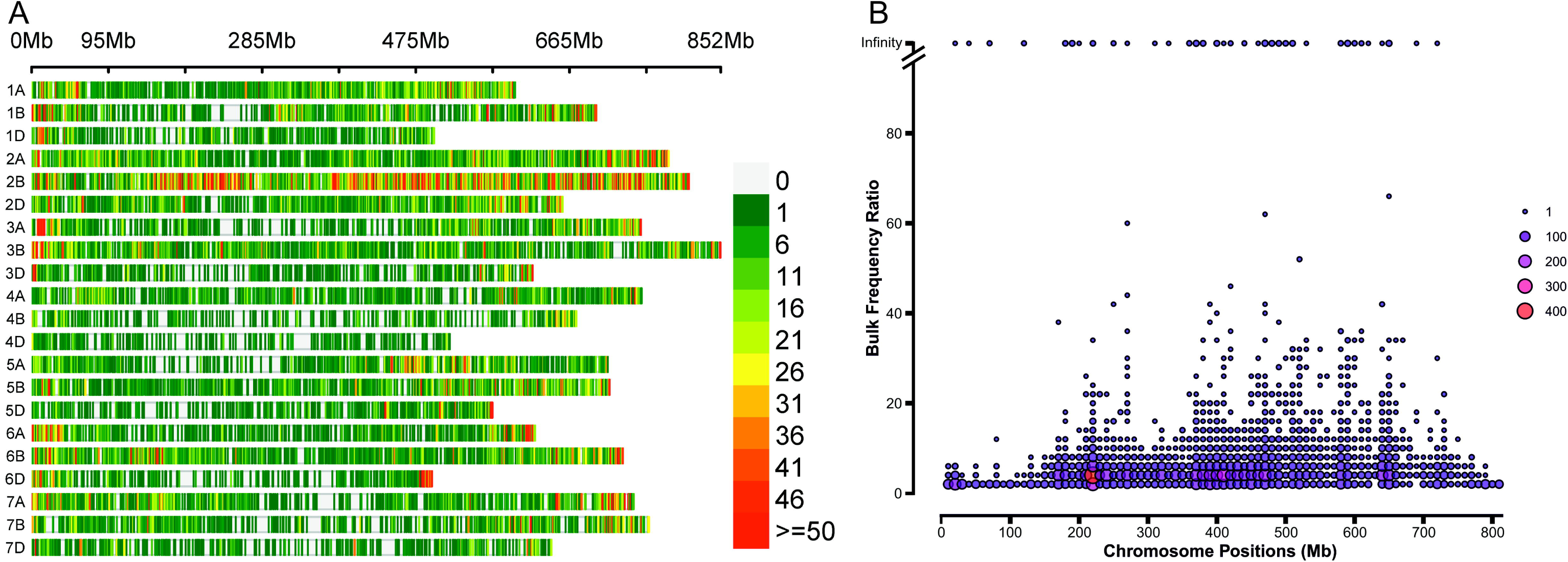
Chromosome-wise distribution of SNPs. **A)** Chromosome-wise SNP density plot depicting the number of SNPs within a 1 Mb window size. **B)** The scatter plot displaying the BFR values.

**Table 2.** Number of reads generated and aligned, and number of SNPs identified after filtering in samples from parents (WL711-2U/2B and WL711), and bulks (resistant bulk, RB and susceptible bulk, SB) belonging to the F_2_ population.

| Sample | Total reads | Aligned reads | Alignment % | Number of SNPs |
| --- | --- | --- | --- | --- |
| RP1 | 68399366 | 66114827 | 96.66 | 53247 |
| RP2 | 62598101 | 58429067 | 93.34 | 90044 |
| SP1 | 66507494 | 64745045 | 97.35 | 73703 |
| SP2 | 61736700 | 56032229 | 90.76 | 66542 |
| RB1 | 60873158 | 58986090 | 96.90 | 92786 |
| RB2 | 52753499 | 51086488 | 96.84 | 83345 |
| RB3 | 55105620 | 53424899 | 96.95 | 87530 |
| SB1 | 57729151 | 51269259 | 88.81 | 75908 |
| SB2 | 56032476 | 52631305 | 93.93 | 69492 |
| SB3 | 69599370 | 61971279 | 89.04 | 80261 |

(RP1= resistant parent replication 1, RP2= resistant parent replication 2, SP1= susceptible parent replication 1, SP2= susceptible parent replication 2, RB1= resistant bulk replication 1, RB2= resistant bulk replication 2, RB3= resistant bulk replication 3, SB1= susceptible bulk replication 1, SB2= susceptible bulk replication 2, and SB3= susceptible bulk replication 3)

**Table 3.** A comprehensive overview of the distribution of polymorphic SNPs on 21 chromosomes of wheat.

| Chrom | Total SNP | SNP having >15 BFR | Chrom | Total SNP | SNP having >15 BFR | Chrom | Total SNP | SNP having >15 BFR |
| --- | --- | --- | --- | --- | --- | --- | --- | --- |
| 1A | 7648 | 1 | 1B | 5641 | 0 | 1D | 4641 | 10 |
| 2A | 13506 | 199 | 2B | 22393 | 1251 | 2D | 11617 | 278 |
| 3A | 6294 | 1 | 3B | 7035 | 1 | 3D | 5519 | 1 |
| 4A | 4196 | 0 | 4B | 3714 | 0 | 4D | 3482 | 1 |
| 5A | 6241 | 0 | 5B | 5671 | 3 | 5D | 5553 | 3 |
| 6A | 4934 | 0 | 6B | 7692 | 0 | 6D | 4724 | 1 |
| 7A | 7810 | 0 | 7B | 4440 | 0 | 7D | 5265 | 3 |

### Distribution of polymorphic SNPs across wheat chromosomes

The total number of SNPs, distributed across all chromosomes, was 148,019. The subgenome B had the highest number of SNPs (56686), while the subgenome D had the lowest number (40801). The chromosome 2B had the maximum SNP markers (22,393), followed by the chromosomes 2A and 2D, where the number of SNPs were 13,506 and 11,617, respectively. Moreover, chromosome 4D carried the minimum SNPs (3482) (Table 3).

The details of the number of SNPs having 0 to infinite BFR values were represented by the figure and table for RB1 vs. SB1, RB2 vs. SB2, RB3 vs. SB3, and RB vs. SB bulks (Table 2, Fig. 2A, 2B). There were 51794, 70477, 8904, 9797, 2766, and 5102 SNPs in the bulk RB vs. SB that had BFR values between 0 and 1, 1 to 3, 3 to 5, 5 to 10, 10 to 15, and 15 to infinity, in that order (Tables 2, Table S1, Fig. 3).

**Figure 3.**
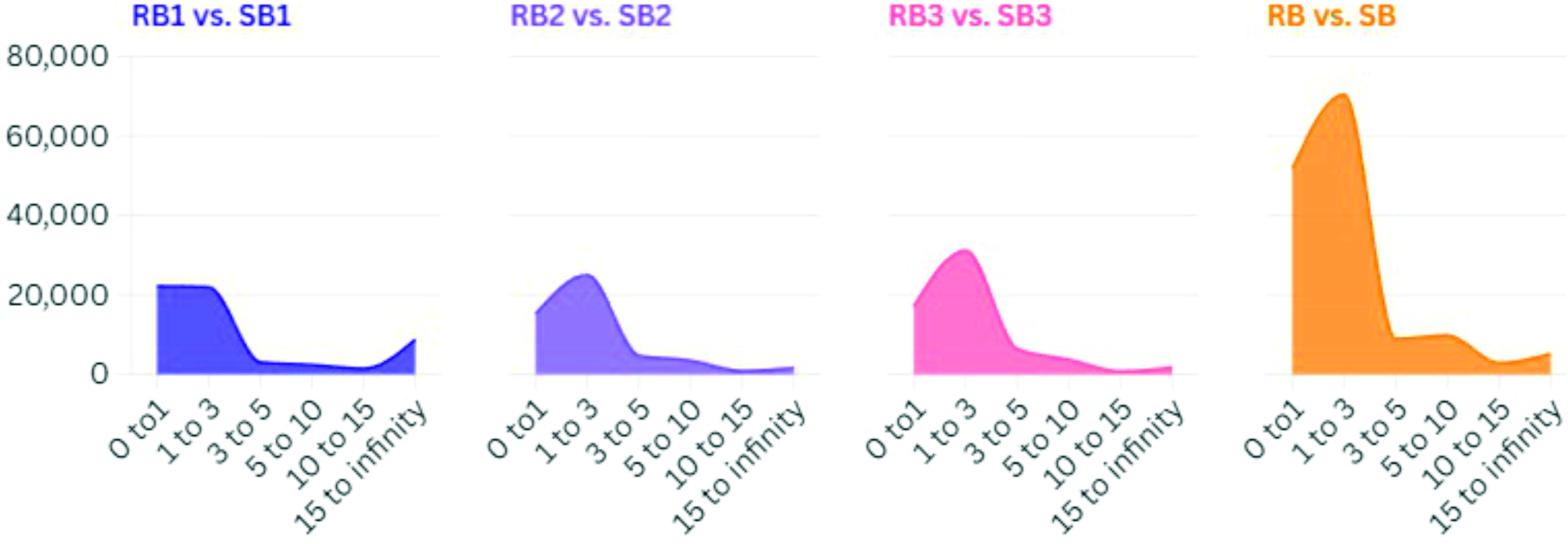
Number of SNPs identified in the samples. The grid of the area chart represents the number of SNPs in 4 bulks with different BFR values

### BFR and polymorphic SNPs

The study identified a total of 1,48,316 SNP counts from the wheat samples used (Table S1). We examined SNPs with > 15 BFR values in the comparison of SB vs. RB, which involved pooling all three resistant bulks and also pooling all three susceptible bulks. Following this process, we identified two bulks: SB and RB. We found 1755 SNPs on 729 transcripts in the comparison of two bulks (SB vs. RB), suggesting their potential association with powdery mildew. Out of all, chromosome 2B exhibited the highest number of SNPs (1,251) having a BFR value of >15 (Fig. 2B, Fig. 3). This was followed by chromosomes 2D and 2A, where the numbers of SNPs were 278 and 199, respectively, with a BFR value of >15. This suggests that chromosome 2B is most likely the carrier of powdery mildew resistance loci.

### Conversion of resistant genes associated SNPs into KASP markers

We selected 14 SNPs on the basis of BFR value > 17 near the flanking region of the QTL (530235618 and 530235656 bp), identified by our QTLseq study (personal communication), and later these were converted into KASP markers. Out of 14, only five markers were validated and utilized in the QTL mapping. In Table S2, highlighted SNPs were validated for QTL mapping.

### Development of high-density map using previously and newly identified molecular markers and QTL mapping

To construct the genetic map, we conducted a parental polymorphic survey using 25 SSR markers positioned on chromosome 2B. The survey revealed that out of these markers, only three displayed polymorphisms. Subsequently, we incorporated the seven SNP markers (out of seven two from QTLseq study; *PmR530235618_2B* and *PmR530235656_2B*) to create the genetic linkage map. We used total 10 markers to screen 114 samples, which included two parents, two bulks, and 110 plants of F2 populations. Only five SNPs among these nine markers successfully underwent validation along with the three polymorphic SSR markers (*Xbarc349*, *Xgwm526*, and *wmc627*) and an additional two markers validated in the QTLseq experiment within the F_2_ segregating population, as depicted in Fig. 4. The present study identified a major QTL (it behaves like a major gene and may be tentatively designated as *PmAT*) with a LOD of 70.3 and a substantial PVE of over 64.7%, confirming the major effect (Table 4). Fig. 5 and Table 3 provide the genomic location of the novel powdery mildew-resistant QTL. The identified gene is located within the position of two SNP markers, i.e. *PmR531114848_2B* and *PmR530235618_2B*, in which a marker (*PmR530235618_2B*) has already been identified in the QTLseq experiment. The distance between these two markers is 0.87 cM, indicating their close proximity. Additionally, we estimated the additive and dominant effects of QTL, finding that the effects of both were -3.9 (suggesting the role in significant reduction in disease severity therby enhaced resistance) and -3.7(indicating the F2 heterozygotes were also showing resistance similar to resistant homozygote alleles indicating their role in resistance), respectively.

**Figure 4.**
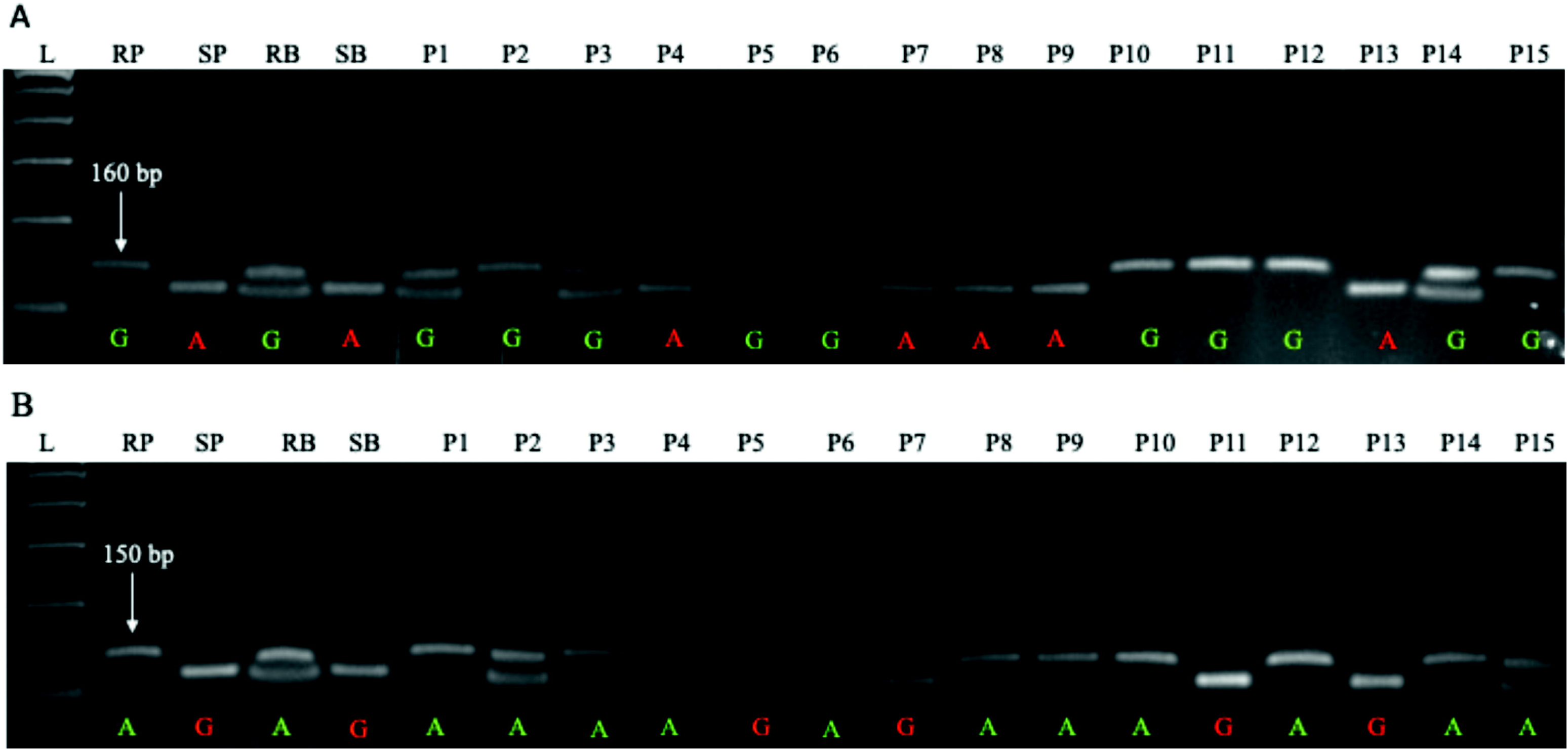
Validation of KASP markers *-* Representative figures of two KASP markers (*PmR531114848*_2B and *PmR530235618*_2B) tightly linked with the powdery mildew-resistant gene (*PmAT*) identified in the present study. **A)** The representative figure of the flanking marker *(PmR531114848_*2B) identified in the present study **B)** The representative figure of the flanking marker (*PmR*530235618_2B) was identified in our previously conducted QTLseq study (unpublished data). This marker was also utilized in the current study during the genotyping of 114 plants in the F2 population for QTL mapping. Whereas, RP = resistant parent; SP = susceptible parent; RB = resistant bulk; SB = susceptible bulk, and P1 to P15 = representative set of 15 individual plants of F_2_ mapping population (n=110). The SNP bases are indicated at the bottom of the gel images.

**Figure 5.**
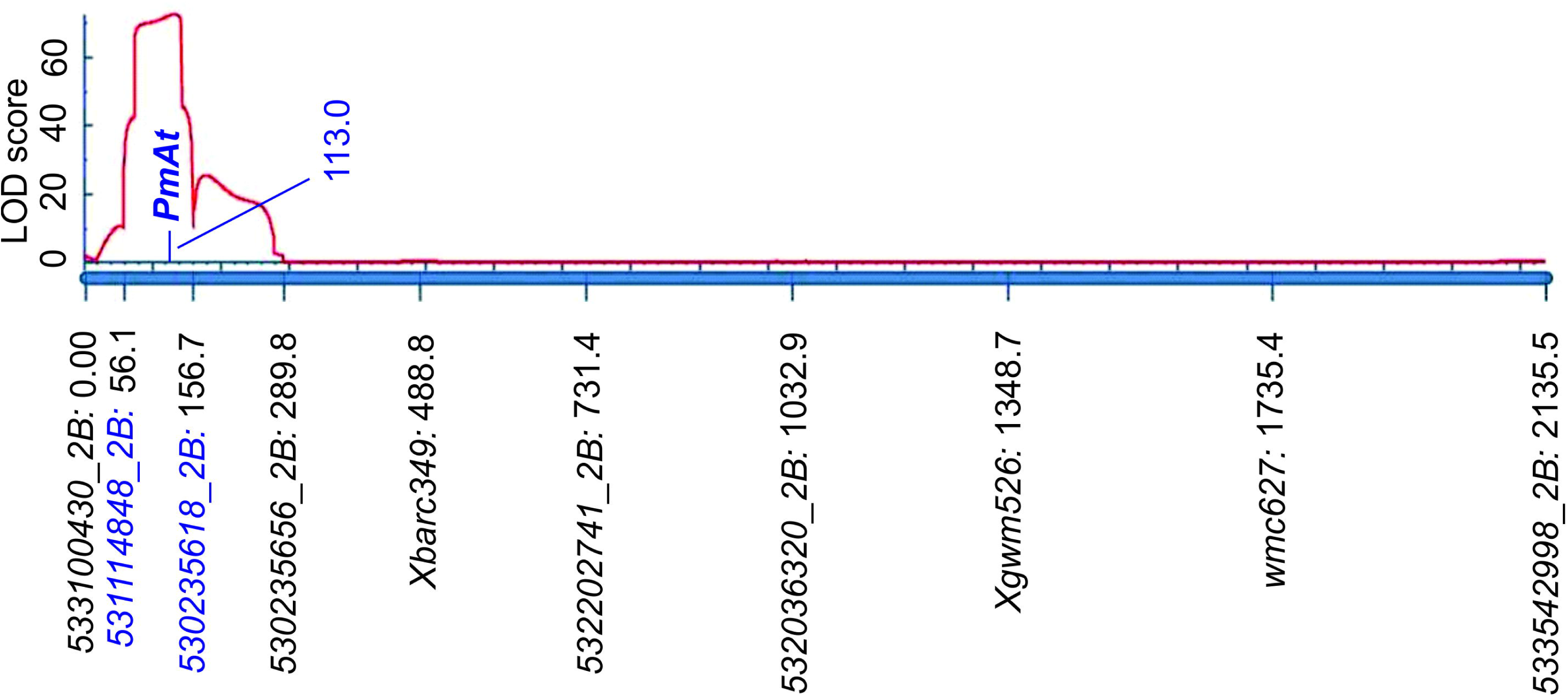
Genetic linkage map developed for Powdery Mildew resistance. The map explains the position of QTL along with the associated markers and LOD score.

**Table 4.** QTL characterization for powdery mildew resistance in F_2_ population of wheat.

| QTL/Gen e | Position (cM) | Left Marker | Right Marker | LO D | PVE (%) | Ad d | Do m |
| --- | --- | --- | --- | --- | --- | --- | --- |
| *PmAT* | 131 | *PmR531114848_2*<br>*B* | *PmR530235618_2*<br>*B* | 72.4 | 64.7 | -3.9 | -3.7 |

### Identification of positional candidate genes in the region linked with powdery mildew resistance

We analyzed interval regions of the identified QTL to extract highly significant and annotated candidate genes. In this study, we found five candidate genes in the QTL interval region (531114848 bp to 530235618 bp). The five candidate genes encode proteins associated with the pentatricopeptide repeat, B-box-type zinc finger, P-loop-containing nucleoside triphosphate hydrolase, and plant peroxidase, among others (Table 5).

**Table 5.**
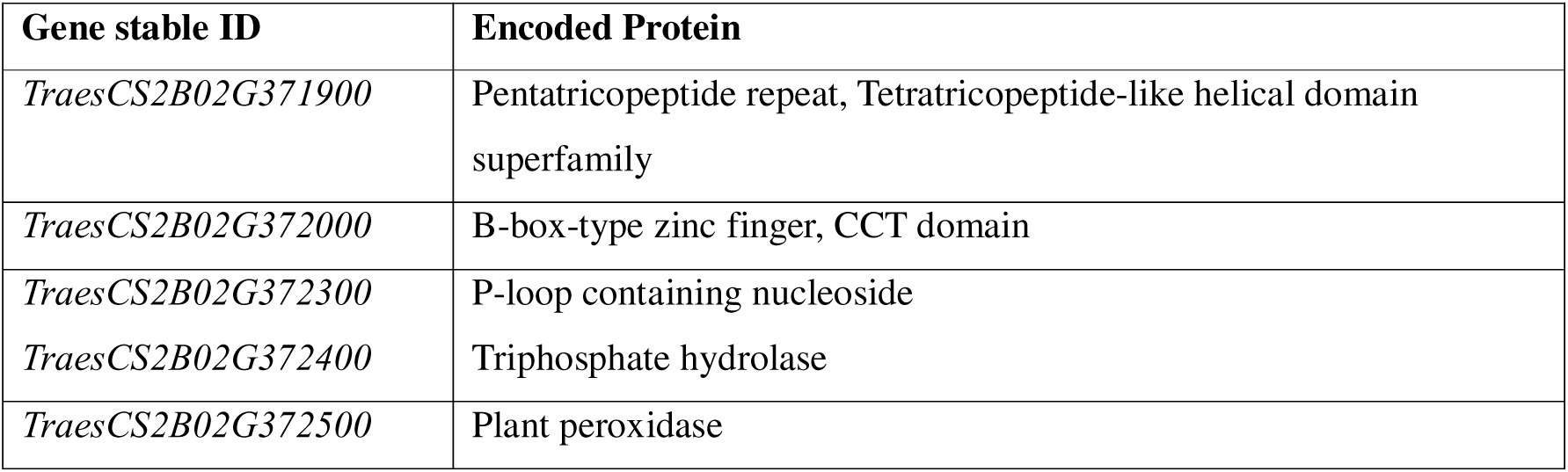
Candidate genes and their encoded proteins found within the interval QTLs.

## Discussion

India’s second most important cereal crop wheat is accounts for one-third of the nation’s grain production and India is only suppressed by China. The biotic and abiotic stresses have major impact on wheat production. To ensure food security for the global population, wheat must be safeguarded from a variety of biotic stresses. Powdery mildew accounts for a large fraction of the biotic factors that could damage the wheat crop. Powdery mildew causes major loss to wheat production in India.

In order to find the novel source of powdery mildew resistance, field experiments were carried using F_2_ generation of a cross between WL711-2U/2B (resistance) as female and WL711 (susceptible) genotypes of wheat. The distribution of adult plant resistance (APR) revealed the presence of major gene for resistance with both additive and dominant effects. Previous studies have also reported the coexistence of qualitative and quantitative resistance in modern wheat cultivars, highlighting the effectiveness and environmental stability of quantitative PM resistance (Keller et al., 1999; Miedaner and Flath, 2007). QTLs, which exhibit additive effects, are known to confer more durable and stable resistance compared to race-specific resistance genes. Consequently, practical breeding programs typically prioritize the selection of cultivars with quantitative resistance (Miedaner and Flath, 2007; Pilet-Nayel et al., 2017).

The *A. triuncialis* translocation happened in the farthest part of WL711’s chromosome arm 2BL and was named T2BS·2BL-2tL (0.95) (Kuraparthy et al., 2007). In this region, a major leaf rust-resistant gene (*Lr58*) has also been identified (Kuraparthy et al., 2007). This introgression line was later tested continuously for more than 15 years for powdery mildew disease resistance, and it was found that the introgression line was also immune against the powdery mildew disease. A disomic substitution line, 5U-5A (BTC17), was also developed using the same parental line, such as WL711 and *Ae. triuncialis*, which harbors gene(s) for powdery mildew resistance on the alien chromosome 5U (Singh et al., 2000; Kamboj et al., 2020). Both the introgression lines are sister lines, but they differ due to the distinct introgression regions for alien chromosomes.

However, researchers have also used other wild relatives of wheat to identify major powdery mildew-resistant genes. For instance, *A. squarrosa* for *Pm2* and *Pm19* (Lutz et al., 1995), *Ae. Longissima* for *Pm66* (Li et al., 2020), *Ae. Searsii* for *Pm57* (Liu et al., 2017), *Ae. Spelltoides* for *Pm32* (Hsam et al., 2003), *Ae. Tauschii* for *Pm34*, *Pm35* and *Pm48* (Miranda et al., 2006, 2007; Fu et al., 2017), *Ae. longissima* for *Pm13* (Cenci et al., 1999), *Elytrigia intermedium* for *Pm40* (Luo et al., 2009), *Haynaldia villosa* for *Pm21* (Lili et al., 1995; Qi et al., 1996), *Secale cereal* for *Pm8*, *Pm17* and *Pm20* (Driscoll and Anderson, 1967; Zeller and Fuchs, 1983; Heun et al., 1990; Friebe et al., 1994), *T. monococcum* for *Pm25* (Shi et al., 1998), *T. sphaerococcum* for *Pm5c* (Hsam et al., 2001), *T. timopheevii* for *Pm27* and *Pm37* (Järve et al., 2000; Perugini et al., 2008), *T. turgidum* for *Pm26*, *Pm30*, *Pm31*, *Pm33*, *Pm36*, *Pm41*, *Pm42*, *Pm68*, *Pm69*, *MG5323* and *PmG3M* (Rong et al., 2000; Hua et al., 2009; Liu et al. 2012; Qiu et al., 2021 Maxwell et al., 2010; Mohler et al., 2005; Zhang et al., 2019; Geng et al., 2016; Yin et al. 2021; Reader and Miller, 1991; Liu et al., 2002; Chen et al., 2005; Blanco et al., 2008; Zhang et al., 2010; Xie et al. 2012; Ben-David et al., 2010; Ji et al., 2008; Ouyang et al.; 2014; Wu et al., 2021; Li et al. 2020), *T. urartu* for *Pm60* (Zhao et al., 2020), *Th. intermedium* for *Pm43* (He et al., 2009) and *Th. ponticum* for *Pm51* (Zhan et al., 2014) were utilised. According to the study of the major resistant genes, we observed that *T. turgidum* was used to find a greater number of resistant genes against powdery mildew in wheat, while *Ae. triuncialis* had not been used before this study to find major powdery mildew-resistant genes.

Recently, a protocol combining bulked segregant analysis (pooling DNA) and local QTL mapping via KASP genotyping was developed to rapidly map major genes in an F_2:3_ population (Hu et al., 2019; Zhan et al., 2021). The aim of this research entailed using BSR-Seq to identify the location of powdery mildew resistance gene *PmAT* before employing KASP markers to precisely locate this gene in an extensive FL population.For the mapping of QTL related to powdery mildew resistance, we have also validated 25 SSR markers on chromosome 2B for QTL mapping and only three were found polymorphic. In this study, only linkage was found between *PmR531114848_2B* and *PmR530235618_2B* with resistant. More interestingly, *PmR*531114848 marker was showing a tight linkage with the QTL. This QTL alone provides more than 64.7% PVE with the LOD value 72.4. The result of this study proves the novelty of the newly identified gene with major effect.

SNP marker (*PmR530235618_2B*) has already been identified in the QTLseq experiment (personal communication), having a 0.87 cM distance and also exhibiting a noteworthy additive effect of -3.9 and a substantial dominant effect of -3.7. The significant additive effect suggests a consistent and measurable enhancement of resistance associated with specific alleles, while the pronounced dominant effect underscores the importance of heterozygosity in conferring additional resistance. Due to the high LOD value (72.4) and PVE% (64.7), this QTL behaves like a major gene. This ensures that the newly discovered gene has a unique location. Several crop species, including wheat, used the BSR-Seq approach to identify QTLs or genes for different traits (Wang et al., 2017; Hao et al., 2019; Zhan et al., 2021; Ma et al., 2021; Qian et al., 2024; Yu et al., 2024). The R gene *PmAT* was initially shown to be associated with the neighboring candidate interval 530.2–531.1 Mb in chromosomal arm 2B using BSR-Seq. Before *PmAT* was found, 11 Pm genes on chromosome arm 2BL had been identified: *Pm6* (Wan et al., 2020), *Pm33* (Zhu et al., 2005), *Pm51* (Zhan et al., 2014), *Pm52* (Wu et al., 2019), *Pm63* (Tan et al., 2019), *Pm64* (Zhang et al., 2019), *PmQ* (Li Y. et al., 2020), *PmKN0816* (Wang et al., 2021), *MlZec1* (Mohler et al., 2005), *MlAB10* (Maxwell et al., 2010), and *PmYD588* (Ma et al., 2021). *PmAT* is different from *Pm6* (698.3–699.2 Mb), *Pm33* (779.1–784.3 Mb), *Pm51* (709.8– 739.4 Mb), *Pm52* (581.0–585.0 Mb), *Pm63* (710.3–723.4 Mb), *Pm64* (699.2–705.5 Mb), *PmQ* (710.7–715.0 Mb), *PmKN0816* (700.4–710.3 Mb), *MlZec1* (796.7–780.0 Mb), *MlAB10* (796.7–780.0 Mb), and *PmYD588* (453.7–506.3 Mb and 584.1–664.2 Mb). This shows that *PmAT* is most likely a new powdery mildew resistant gene. However, further evidence is required to demonstrate the connection between these powdery mildew resistant genes on chromosomal arm 2BL. Mutual allelism tests and even cloning these genes are two examples of how to accomplish this.

The candidate gene *TraesCS2B02G371900* encodes a protein with two key domains, Pentatricopeptide repeat (PPR) and Tetratricopeptide-like helical domain superfamily. Pentatricopeptide repeat-containing proteins are crucial for RNA-binding proteins post-transcriptional processes within the mitochondria and chloroplasts (Delannoy et al., 2007). These proteins are involved in several physiological roles, including defense against necrotrophic fungi (Laluk et al., 2011). Pentatricopeptide repeat (PPR) proteins represent a major locus utilized in hybrid wheat breeding programs (Walkowiak et al., 2020). Pentatricopeptide repeat (PPR) and Tetratricopeptide-like helical domain superfamily are also involved in the defence response of wheat against leaf rust (Vikas et al., 2022) and stripe rust pathogens (Singh, 2022).

The potential gene *TraesCS2B02G372000* encodes one or two B-BOX domain structures have zinc finger transcription factors that belong to the B-BOX domain protein family (Gangappa and Botto 2014). According to numerous studies, BBX proteins play a role in several activities, such as signal transduction, flower development, shade avoidance, and plant photomorphogenesis (Azam et al., 2024; Gangappa et al., 2013). Interestingly, recent research has suggested that BBX proteins contribute to plants’ ability to withstand infections. In Oryza sativa, overexpressing the BBX protein gene OsCOL9 boosts resistance to blast disease, whereas knocking out the gene increases susceptibility to the disease. The ethylene (ET) and salicylic acid (SA) signaling pathways are linked to this resistance (Liu et al., 2016). IbBBX24 overexpression increases *Ipomoea batatas’* resistance to *Fusarium graminearum*-induced wilt disease, but IbBBX24 silencing decreases such resistance. The JA signaling pathway is linked to this resistance (Zhang et al., 2024). The role of this protein has also been observed in wheat against the powdery mildew pathogen and found downregulation in the genes that encode B-BOX protein (Vishwakarma et al., 2023).

*TraesCS2B02G372300* and *TraesCS2B02G372400* are candidate genes that encode proteins P-loop containing nucleoside and Triphosphate hydrolase involved in defense response against a range of diseases. A possible involvement for P-loop protein in disease resistance has been suggested by their association with plant defense responses (Juliana et al., 2018; Qi et al., 2024; Salam et al., 2023). The role of P-loop containing nucleoside protein has been observed in resistance against leaf rust (Juliana et al., 2018), stripe rust (Singh, 2021) and spot blotch (Kaur et al., 2023) disease in wheat. A gene in rice that encodes a loop-containing nucleotide triphosphate hydrolases superfamily protein was found to protect the plant from blast disease (Zheng et al. 2004).

The candidate gene *TraesCS2B02G372500* encodes for plant peroxidase, because they support the plant’s structural and biochemical defenses against infections, peroxidases are an essential part of its defensive mechanism. Peroxidases that are also part of the PR-9 family of pathogenesis-related proteins play an important role in defense response against diseases in plants (Dos Santos and Franco, 2023). The production of reactive oxygen species (ROS) during plant-pathogen interactions is one of their main roles (KámánLTóth et al., 2019). This oxidative burst serves as a primary defense mechanism by limiting the entry of pathogens into the plants (KámánLTóth et al., 2019). The upregulation in the genes related to the peroxidases in wheat against powdery mildew have also been reported (Liu et al., 2005; Mustafa et al., 2016)

## Conclusion

The research adds to powdery mildew wheat defense through the discovery and evaluation of the new resistance gene *PmAT* introgressed from *Ae. triuncialis*. Wild wheat relatives gain significance as sources to find new resistance genes through the combination of BSR-Seq sequencing technology to improve wheat breeding programs. Analysis of 148,000 SNPs revealed that chromosome 2B contained the most high-BFR SNPs out of all regions and showed the strongest association with resistance. The fourteen highest priority SNPs were transformed into KASP markers to establish a genetic linkage map alongside five verified markers. This marked region contained five genes that control defense processing and stress communication pathways. Further, this should proceed with the cloning of this gene because researchers have not identified the segments translocated from *Ae. triuncialis*. Also, the exact molecular processes associated with the *PmAT-* mediated powdery mildew resistance will be useful in developing improved resistant wheat cultivars.

## Data availability

All data are available from the corresponding authors upon reasonable request. Transcriptomic data were deposited in the NCBI database under BioProject accession number: PRJNA1249271.

## CRediT authorship contribution statement

Ramandeep Kaur: Writing - Original draft, Investigation, Formal analysis. Raman Dhariwal: Formal analysis, Writing - Review & Editing. Imran Sheikh, Harcharan S. Dhaliwal, M. Sivasamy, Thamaraikannan Sivakumar, Sundeep Kumar: Writing - Review & Editing. Neeraj Kumar Vasistha: Writing – Investigation & Supervising, Review & Editing, Funding acquisition, Conceptualization.

## Acknowledgments

We express our appreciation to Science and Engineering Research Board (SERB), India for supporting NKV with a Start-Up Research Grant (SRG/2020/000091) and the Department of Genetics-Plant Breeding and Biotechnology, Dr. Khem Singh Gill Akal College of Agriculture, Eternal University, Baru Sahib, Sirmour, India, for providing facilities to Neeraj Kumar Vasistha, and Ramandeep Kaur to undertake this research. The helpful assistance from ICAR - IARI Regional Station, Wellington, in taking off-season crops is gratefully acknowledged.

